# Multilevel modeling of cell population migration cycles and response to environmental pH in *Bacillus subtilis*

**DOI:** 10.1101/2020.09.11.292474

**Authors:** Madoka Nakayama, Izumi Takagi, Jun-ichi Wakita, Wataru Shoji, Sohei Tasaki

## Abstract

Microbial populations are ubiquitous, many of which form robust, multicellular structures called biofilms that protect cells from environmental damage. Understanding their migration cycles is crucial for controlling target microbial populations. Here we develop a multilevel mathematical model that captures the migration cycle of *Bacillus subtilis* cell populations and reproduces the periodic expansion of colonies into concentric circles. We construct an input/output model that regulates cell types in response to environmental conditions; based on this, we propose a theoretical model for the generation of cell population migration cycles. We then construct a multilevel system that links the cell-level state regulation model with tissue-level partial differential equations governing cell density, signal, and resource fields. Through this approach, we successfully simulate cyclically expanding colonies as well as their environmental pH dependence.

The proposed models provide a robust basis for describing highly self-regulating multicellular systems and predicting broader biofilm-related phenomena.

## Introduction

Microbial populations often form colonies and exist stably within robust structures known as biofilms [1–9]. The structural robustness of biofilms makes it difficult to eradicate disease-related microorganisms [10–12]. Conversely, as numerous beneficial microorganisms continue to be discovered, selectively controlling target populations has become an urgent issue [13–15]. The process of biofilm formation involves [1, 2]: (i) bacterial attachment to suitable surfaces; (ii) proliferation and secretion of extracellular polymeric substances; (iii) the eventual dispersal of mobile cells from the enlarged colony to seek new environments. These released cells colonize new sites that satisfy their growth requirements, effectively returning to the initial phase. To summarize, these cells return to the first phase of biofilm formation as described in (i) above. This sequence is known as the life cycle of a bacterial biofilm. Biofilms are complex communities composed of diverse cells; thus, their formation and stabilization involve multilevel interactions, such as information exchange within and among populations. Due to this complexity, the detailed mechanisms underlying their life cycles have not yet been fully elucidated. We aim to develop a multilevel computational model that predicts these migration cycles, thereby contributing to the regulation of biofilm life cycles.

Concentric colony formation serves as a simplified model illustrating the migration cycle observed in biofilms (Fig. 1). *Proteus mirabilis* [16–19] and *Bacillus subtilis* [19–24] are well known for forming such patterns, where the dominant cell population periodically switches between migrating and non-migrating states. In *B. subtilis*, migration is driven by swarming—a collective movement of motile cells [25–28]. Conversely, the non-migrating state is dominated by matrix-producers [25]. There are three possible factors that cause a migrating cell population to enter a non-migrating state: (i) switching to a non-motile cell type; (ii) cessation of hyperflagellation; and (iii) insufficient concentration of surface-wetting molecules, such as surfactin. In the case of this *B. subtilis* concentric colony, the non-migrating cell population is in the form of chains of proliferating matrix producers, and the cells do not move at all [22]. Therefore, the major mechanism underlying the stoppage of swarming involves the predominance of non-motile cell types, most of which are matrix producers. Although the regulation of surface-wetting molecule secretion and hyperflagellation should also be involved in the stabilization of the cycle [22], we focus on cell-type switching as the primary source of cycle generation.

**Fig. 1:**
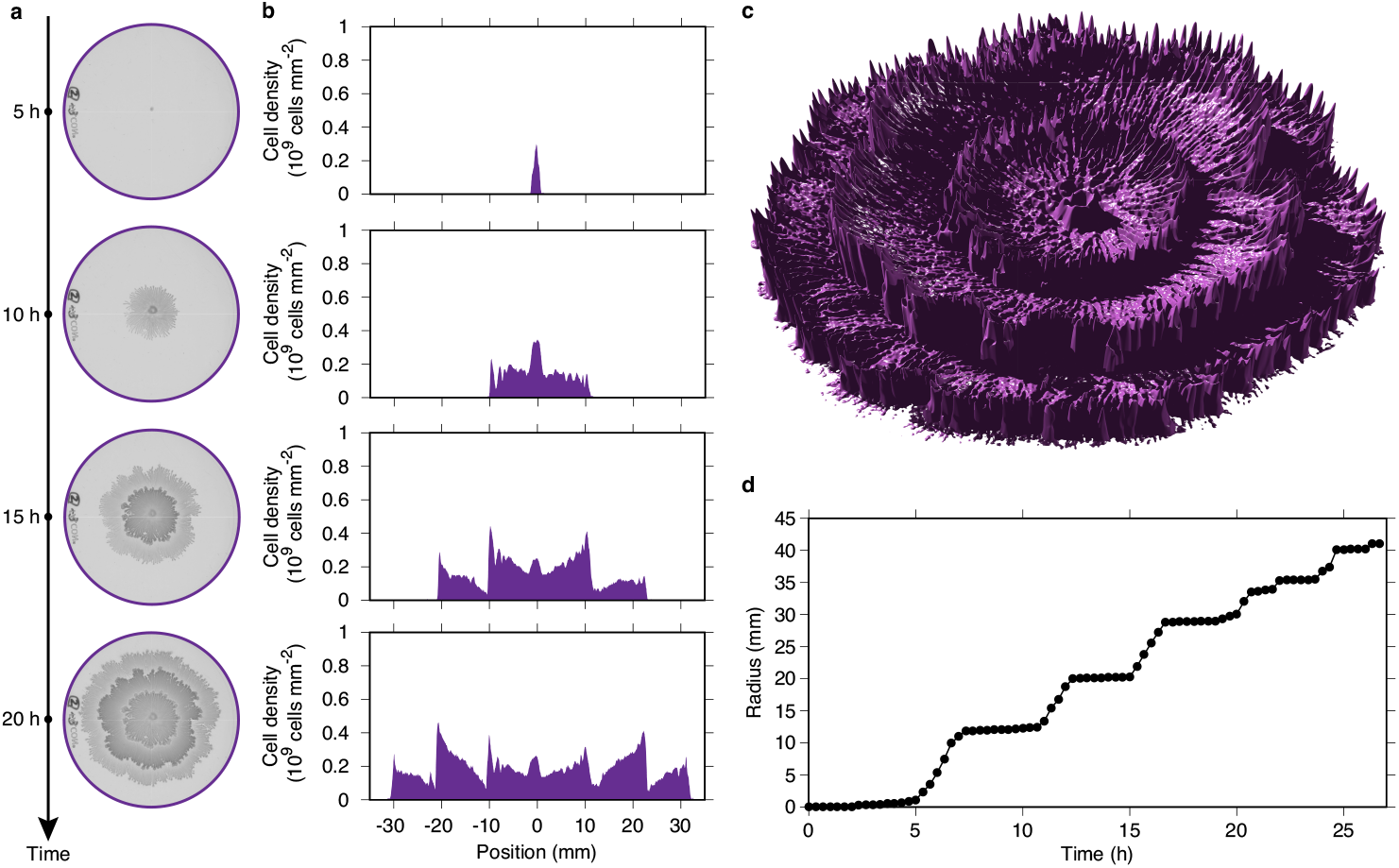
Formation of concentric colonies. **a**, Growth of concentric colonies. The center is the inoculation point. The diameter of the dish is 88 mm. Incubation time, 5, 10, 15, and 20 h after inoculation. **b**, Cross-section cell density. The center is the inoculation point. **c**, Three-dimensional reconstruction of the colony. **d**, Increasing colony radius.

Swarming by motile cells drives the migration phase, causing the colony to rapidly spread across the medium surface in two dimensions. The colony is extremely thin, consisting of almost a single layer of cells. In contrast, matrix producers do not move, and proliferate while producing extracellular matrices [29]. The colony not only expands two dimensionally but also increases in thickness (grows in the direction perpendicular to the medium plane). Cells called surfactin producers [30, 31] are known to coexist to assist the slow two-dimensional expansion in this phase, improving sliding mobility on the medium surface [32], although it is far behind the expansion speed during the migration phase. Thus, two distinct phases alternate: the “migration phase,” during which motile cells rapidly expand the colony area, and the “growth phase,” during which non-motile cells increase the thickness of the colony. This cycle constitutes the migration cycle of a *B. subtilis* cell population.

Here, we construct a multilevel mathematical model to describe the dynamics of spatio-temporally heterogeneous cell populations and predict their response to environmental changes. We specifically investigate the effect of environmental pH on cell-type selection and concentric colony formation. By combining a cell-type regulation model based on gene networks with tissue-level equations, we provide a plausible explanation for the formation of concentric patterns. Our model offers a new framework for the predictive modeling of multicellular systems responding to environmental fluctuations.

## Results

### Concentric colonies embody the cell population migration cycle

We prepared a nutrient-rich agar medium with a concentration of 0.7%, representing an intermediate stiffness within the typical range for solid agar media, and inoculated *B. subtilis* at the center of the plate. When cultured at approximately 35°C, the bacteria formed concentric colonies on the medium surface (Fig. 1a–c, Extended Data Fig. 1). The spatio-temporal periodicity observed in the growth of these colonies directly reflects the cell migration cycle. By examining the leading edge of the colony under a microscope (Extended Data Fig. 2) and monitoring the expansion rate of the colony radius (Fig. 1d), we were able to clearly distinguish between the migration phase and the growth phase. During a single cycle—comprising one migration phase and one growth phase—the rate of increase in the total cell number remains relatively constant. However, upon entering a new cycle and transitioning into motile cells, the cell proliferation rate undergoes a temporary decrease. This reduction potentially reflects the high metabolic cost and nutrient consumption associated with the differentiation into and maintenance of the motile cell state (Extended Data Fig. 3).

### Conditions for the emergence of cell population migration cycles

The specific conditions for the formation of concentric *B. subtilis* colonies have been a subject of long-standing investigation. Previous studies reported that within an optimal temperature range of 35–37°C, even under favorable nutritional conditions, concentric colonies form only within an extremely narrow range of agar concentrations (0.65–0.68%) [22]. This suggests that additional stress parameters, such as low temperature [20], are required to stabilize these patterns. In this study, we hypothesized that environmental pH acts as a critical stress parameter. To test this, we fixed all other conditions—including agar concentration at an intermediate level of 0.7%— and systematically varied the environmental pH (Fig. 2). In the neutral pH range, no periodic growth was observed; matrix producers remained the predominant cell type, and colonies expanded primarily by increasing their volume through proliferation (Fig. 2a,b). As the environmental pH decreased, stochastic migration phases began to emerge around pH 6.8. Near this transition point, spatial variation was highly pronounced, periodicity remained unclear, and the resulting patterns deviated significantly from concentric circles. However, when the pH dropped below 6.5, stable, periodically expanding concentric colonies were successfully formed. Conversely, at pH values below 5.3, colony development became erratic, often stopping prematurely. Finally, at pH levels below 5.1, colony formation was completely inhibited. These results identify a stable pH range of 5.3–6.5 for the cyclic growth of concentric colonies.

**Fig. 2:**
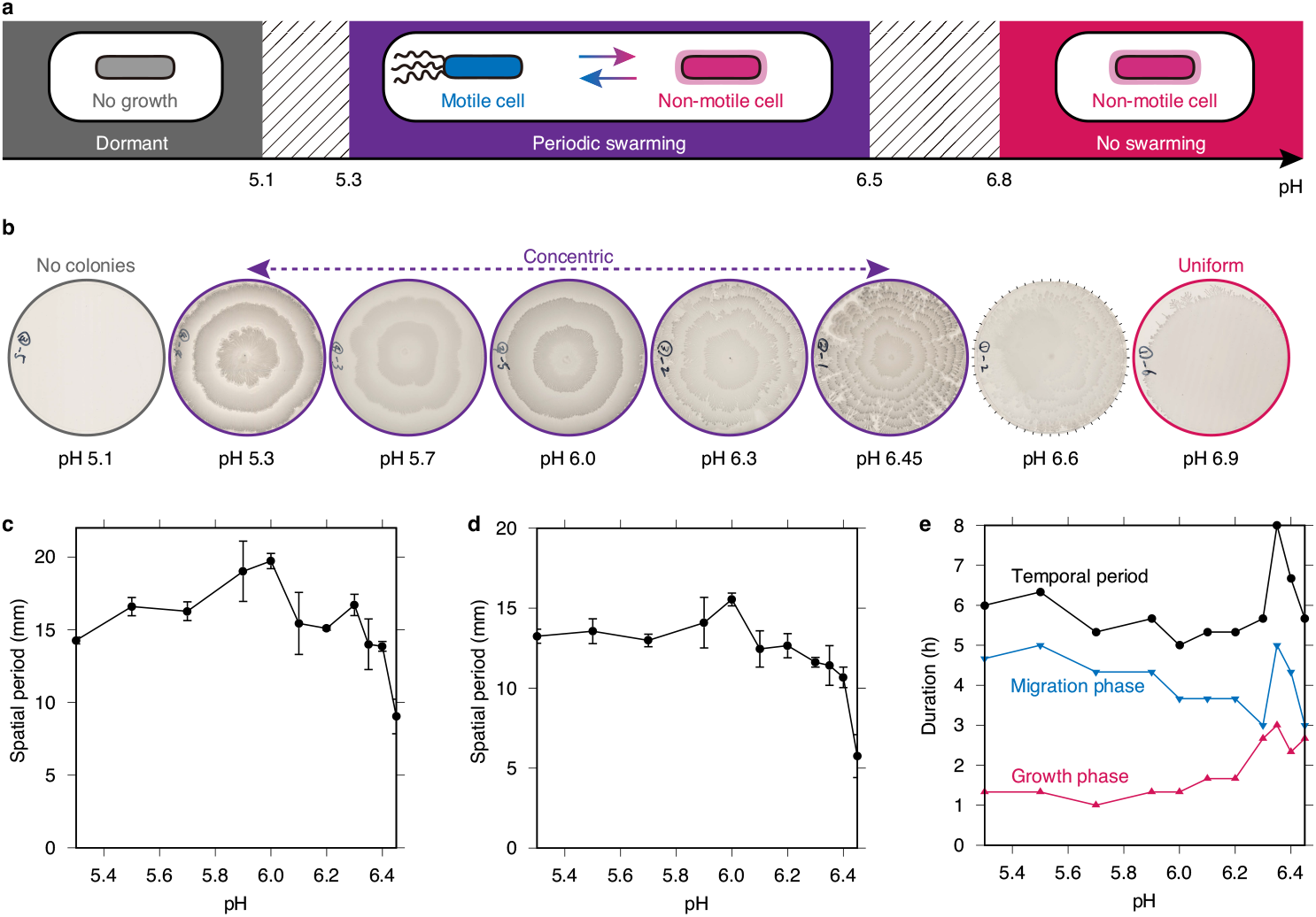
Response of colony patterns to changes in environmental pH. **a**, Dominant cell types that determine colony growth patterns. **b**, Colonies under different environmental pH conditions. **c**,**d**, Spatial period. The depth of the first terrace (**c**), The depth of the second terrace (**d**). **e**, Phase duration: growth phase, migration phase, and their total (migration cycle). Data represent mean ± s.d., *n* = 6 (**c, d**).

Next, we characterized the properties of the concentric colonies within this stable pH range. First, the circularity of the rings remained relatively constant, despite some morphological variation observed near the neutral pH transition (Extended Data Fig. 4). The spatial period (the depth of each terrace) was essentially maintained at a constant value between pH 5.3 and 6.0 (Fig. 2d). However, as the pH increased toward the transition range (pH 6.0–6.5), the spatial periodic structure began to collapse, with the periodicity decreasing sharply just prior to the transition.

To clarify the underlying mechanisms of these morphological shifts, we analyzed the durations of the migration and growth phases (Fig. 2e). Notably, while the total duration of the migration cycle remained nearly constant, the relative length of each phase shifted: the migration phase duration increased as pH decreased, while the growth phase duration decreased accordingly. These findings indicate that while finer spatial patterns reflect shorter migration periods, the overall life cycle of the cell population remains homeostatic, regardless of environmental pH fluctuations.

### Environment-dependent regulation of cells and cell populations

We investigated the mechanisms controlling the generation of and switching between the two distinct phases within the cell population. Coordination at the population level must be understood through cell-level regulation in response to environmental conditions. The transition between motile cells and matrix producers is a well-studied regulatory system, particularly regarding the switch between planktonic and biofilm states [25, 33–36]. These two cell types are characterized by mutually exclusive gene expression states. We developed a simplified mathematical model to explore these cellular state dynamics (Fig. 3a). In this model, cell states are described by four grouped variables representing coordinated genes and their products: (1) *S*: the group represented by phosphorylated Spo0A (Spo0A∼ P), including the phosphorelay components Spo0F and Spo0B; (2) *H*: SigH; (3) *A*: AbrB; (4) *C*: the group represented by ComK, driven by the ComX-ComP-ComA pathway.

**Fig. 3:**
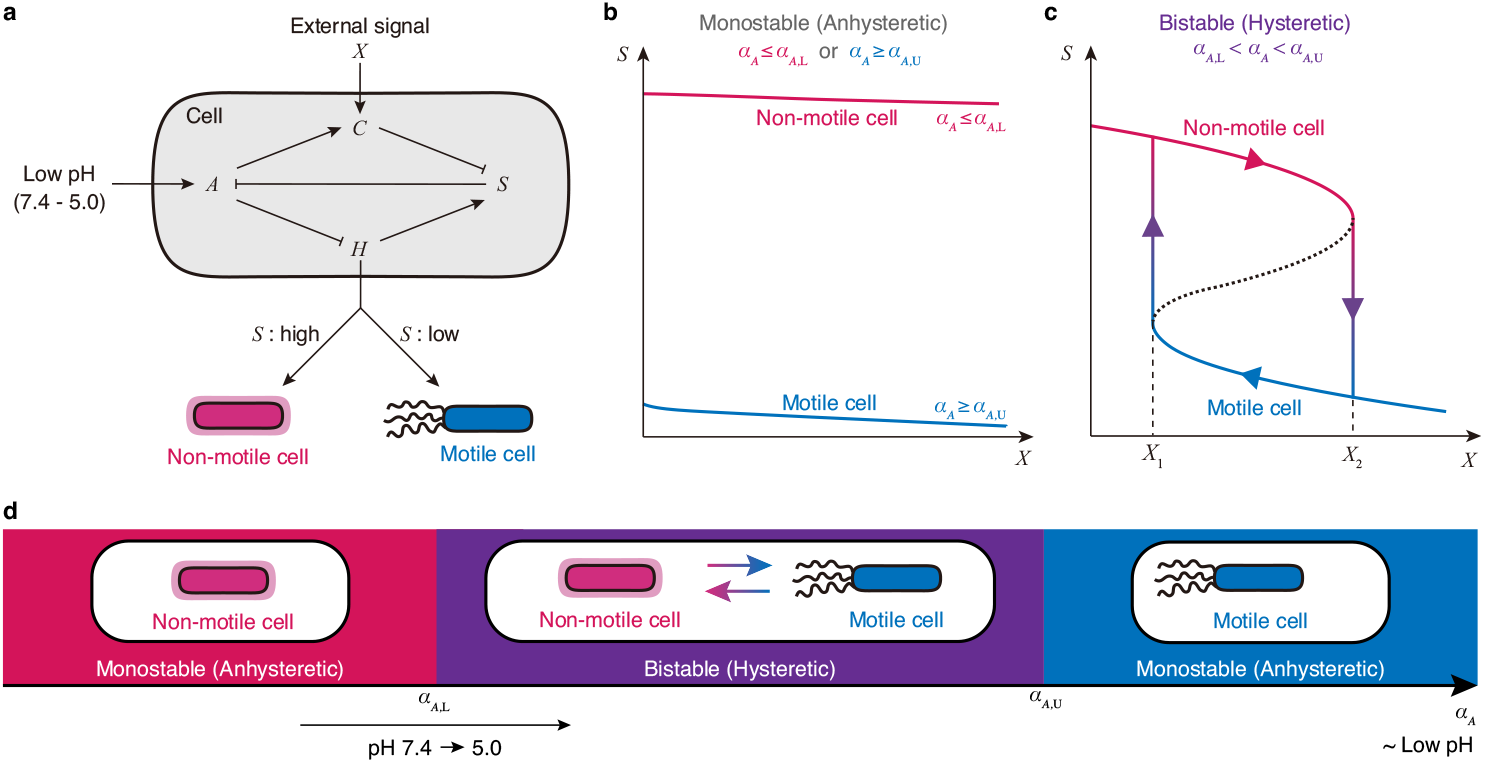
A model for the response of cells and cell populations to environmental pH. **a**, A model for cell-type selection for environmental pH and cell density. **b**,**c**, Two types of cell-type controls: Anhysteretic (**b**) and hysteretic (**c**). **d**, A model for cell population-type selection. The horizontal axis (*α*_*A*_) is a parameter indicating the low pH of the environment.

The output of this control system detemines the cell type: a matrix producer when *S* is high, and a motile cell when *S* is low. While sporulation is initiated when *S* levels remain consistently very high, it typically occurs during late-stage nutrient depletion, long after colonization. Therefore, sporulation was excluded from this model. The inputs to the system include external environmental conditions and auto-inducing signals representing cell density. Given that our experiments used high-nutrient media, the nutritional conditions were assumed to be constant and sufficient during colony formation. For the cell-density signal, we used the concentration of the secreted peptide ComX [37–39], denoted as *X*, which acts as an input from *C* to the cell-type regulation system [37, 38, 40]. A key focus here is how environmental pH integrates into this system. Based on previous studies, *H* or *A* are the primary candidate input points for external pH [41, 42].

Here, we present a minimal, smooth dynamic model of these four variables— *S* = *S*(*t*), *H* = *H*(*t*), *A* = *A*(*t*) and *C* = *C*(*t*)—defined by the following differential equations:

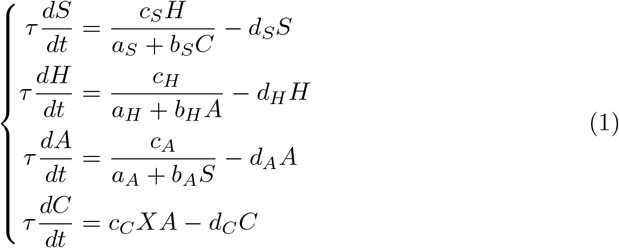

for *t* > 0, the time variable. *τ, a*_*_, *b*_*_, *c*_*_, *d*_*_ (* = *S, H, A, C*) are positive constants; *τ* represents the relaxation time. *a*_*S*_ represents the baseline inhibition rate of *S, b*_*S*_ represents the inhibition rate of *S* by *C*, and *c*_*S*_ represents the activation rate of *S* by *H. a*_*H*_ represents the baseline inhibition rate of *H, b*_*H*_ represents the inhibition rate of *H* by *A*, and *c*_*H*_ represents the baseline activation rate of *H. a*_*A*_ represents the baseline inhibition rate of *A, b*_*A*_ represents the inhibition rate of *A* by *S*, and *c*_*A*_ represents the baseline activation rate of *A. c*_*C*_ represents the expression rate of *C* activated by *d*_*S*_, *d*_*H*_, *d*_*A*_, and *d*_*C*_ represent the decay rates of *S, H, A*, and *C*, respectively.

The curve of the set of equilibrium points of this system can be classified into two types regarding the response to *X*: the variable that represents the cell state (for example, *S*) is monotonic and non-monotonic (Fig. 3b, c) [43]. In the non-monotonic case, the equilibrium curve follows an S-shaped trajectory with two turning points, indicating hysteretic control (Fig. 3c) rather than a simple, anhysteretic switch (Fig. 3b). This hysteretic regulation is essential for the generation of cell population migration cycles and the formation of concentric colonies. When hysteresis occurs in response to fluctuations in *X*, the following migration cycle emerges: Migration phase: As the cell population disperses in the motile state, the local cell-density signal *X* at the growth front decreases. When *X* falls below a specific threshold *X*_1_, the population switches to the matrix-producing state. Growth phase: As the population grows and matures in the matrix-producing state, *X* increases. When *X* exceeds a higher threshold *X*_2_ (> *X*_1_), the population reverts to the motile state.

The overall structure of these equilibrium points in response to fluctuations in *X* in (1) is determined by the environmental parameter *α*_*A*_ (activating *A*) and *α*_*SH*_ (activating *S* and *H*) [43]:

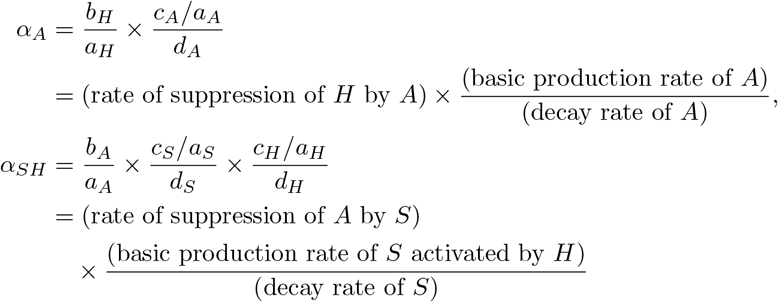

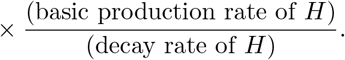

By fixing *α*_*SH*_ and varying *α*_*A*_, the model successfully reproduces the shifts in cell-type control corresponding to changes in environmental pH (Fig. 3d). This theoretical framework explains the mechanism of the migration cycle within the hysteretic parameter range (Fig. 3c) and clarifies its dependence on environmental stress (Fig. 3d).

### Multilevel modeling of colony formation involving dominant cell type transitions

Building upon the theoretical model of population state control, we developed a multilevel model to integrate cellular-level regulation with the tissue-level processes of colony morphogenesis and environmental variation. While agent-based models (ABMs) represent a straightforward approach by coupling individual cell behavior with field equations, they often pose significant challenges due to their high degrees of freedom, the complexity of interactions, and prohibitive computational costs. Consequently, such detailed models may be less effective for uncovering the underlying essence of colony pattern formation. In this study, we aimed to construct a simplified model that captures the core mechanisms of the cell population migration cycle and the resulting patterns. To achieve this simplification, we focused on the observation that a single cell type typically dominates the colony’s leading edge at any given time. Although swarming cells are absent in the inner regions of the colony during the migration phase, these interior cells have negligible influence on pattern formation due to nutrient depletion; therefore, they can be largely ignored in our model. We defined the dominant cell type based on a representative point 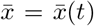 near the colony’s leading edge, where the concentration of the signaling molecule *X* = *X*(*x, t*) is maximal: 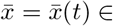 arg max *X*(*x,t*). This point represents the spatial coordinate of the most 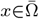 physiologically active cells within the population.

Next, we formulated the tissue-level dynamics within a spatial domain Ω ⊂ ℝ^2^. For the spatial variable *x* ∈ Ω and time *t* > 0, the dynamics of cell density *B* = *B*(*x, t*), nutrient concentration *N* = *N* (*x, t*) and signaling substance *X* = *X*(*x, t*) are governed by the following system of equations:

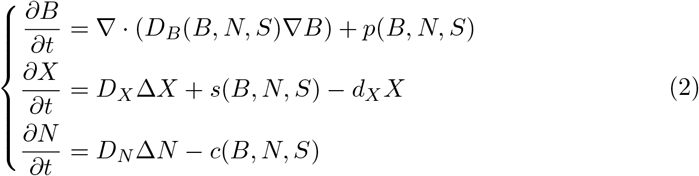

for *x* ∈ Ω and *t* > 0. In this framework, bacteria migrate on the medium surface with a diffusion coefficient *D*_*B*_ and proliferate at a rate *p*. During this process, they secrete *X* at rate *s*, which diffuses through the medium with a diffusion coefficient *D*_*X*_ and undergoes degradation at rate *d*_*X*_. Nutrients *N* diffuse within the medium with 8 a diffusion coefficient *D*_*N*_ and are consumed by the bacteria at a rate *c*. Critically, bacterial motility *D*_*B*_, proliferation *p*, signal secretion *s* and nutrient consumption *c* are functions of local cell density *B*, nutrient availability *N*, and the master regulator *S*: *D*_*B*_ = *D*_*B*_(*B, N, S*), *p* = *p*(*B, N, S*), *s* = *s*(*B, N, S*), *c* = *c*(*B, N, S*).

In summary, the proposed multilevel model links the cellular-state regulatory model (1) with the tissue-level field equations (2) through the representative signal value 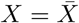, where 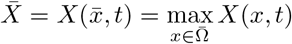.

### Numerical simulation of concentric pattern formation and the migration cycle

Using the proposed multilevel model (1)–(2) with 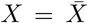, we reproduced the formation of concentric circular patterns that periodically repeat different phases. To achieve this, we selected parameters that generate hysteretic cell-type control (Fig. 3c). For simplicity, we assumed a one-dimensional spatial domain: Ω = (0, *R*) with a system size of *R* = 160. Under appropriate settings (Methods), the model successfully reproduces concentric circular patterns within a specific parameter range (Fig. 4).

**Fig. 4:**
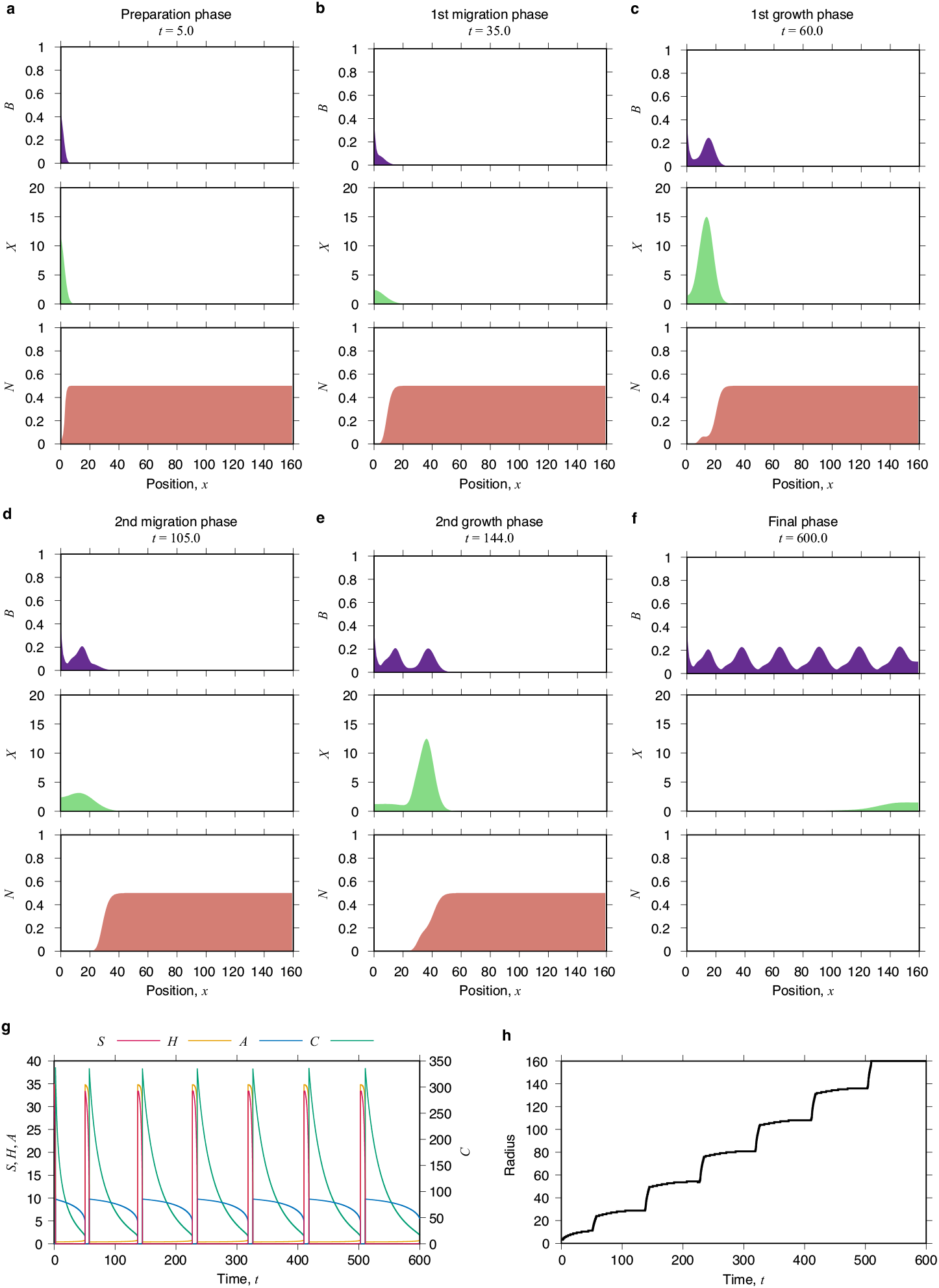
Multilevel simulation of concentric pattern formation. **a**–**f**, Snapshots of *B, X, N*. The preparation phase (0th growth phase) (**a**), the 1st migration phase (**b**), the 1st growth phase (**c**), the 2nd migration phase (**d**), the 2nd growth phase (**e**) and the final phase (**f**). **g**, Time series of *S, H, A, C*. **h**, Colony radius.

The process begins with a preparation phase dominated by matrix producers (Fig. 4a). As these cells proliferate, the concentration of the secreted signal *X* eventually exceeds the threshold *X*_2_ (Fig. 3c), triggering a transition to the motile cell state. This motile population rapidly expands across the culture surface, marking the first migration phase (Fig. 4b). During this collective movement, cell density at the colony’s leading edge gradually decreases, causing the concentration of the signaling molecule *X* to decline. When *X* falls below the lower threshold *X*_1_, the cells revert to the matrix-producing state, initiating the first growth phase (Fig. 4c). Together, these phases constitute the first migration cycle. As the population continues to grow, *X* rises again, eventually surpassing *X*_2_ and triggering the migration phase of the second cycle (Fig. 4d,e). Ultimately, this repeated process results in a periodic spatial distribution of bacteria, while both *X* and *N* approach zero as resources are consumed (Fig. 4f). Throughout this patterning process, the regulators *S* and *H* dominate during the preparation and growth phases, whereas *A* and *C* dominate during the migration phase (Fig. 4g). The expansion rate of the colony radius is rapid during the migration phases and extremely slow during the growth phases (Fig. 4h). The dynamic is highly consistent with our experimental observations (Fig. 1d). However, the temporal change in the total cell number cannot be fully reproduced by this simplified model (Extended Data Fig. 4). This discrepancy arises from the model’s fundamental assumption that all cells at the leading edge synchronize their state transitions, a simplification that overlooks the finer internal population dynamics.

### Simulation of pH-dependent pattern selection and consistency with experiments

We investigate whether our model, (1)–(2) with 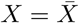, could reproduce the characteristic pattern transitions corresponding to environmental pH fluctuations. As previously discussed, a decrease in environmental pH (within the range of pH 5.0–7.4) is assumed to activate *A*. We therefore mapped *c*_*A*_ to this pH decrease, such that the cellular state control parameter *α*_*A*_ is proportional to *c*_*A*_, while *α*_*SH*_ remains constant. In the following analysis, we varied *h*_*A*_ = 3 −*c*_*A*_ as the primary parameter representing environmental pH (Fig. 5).

**Fig. 5:**
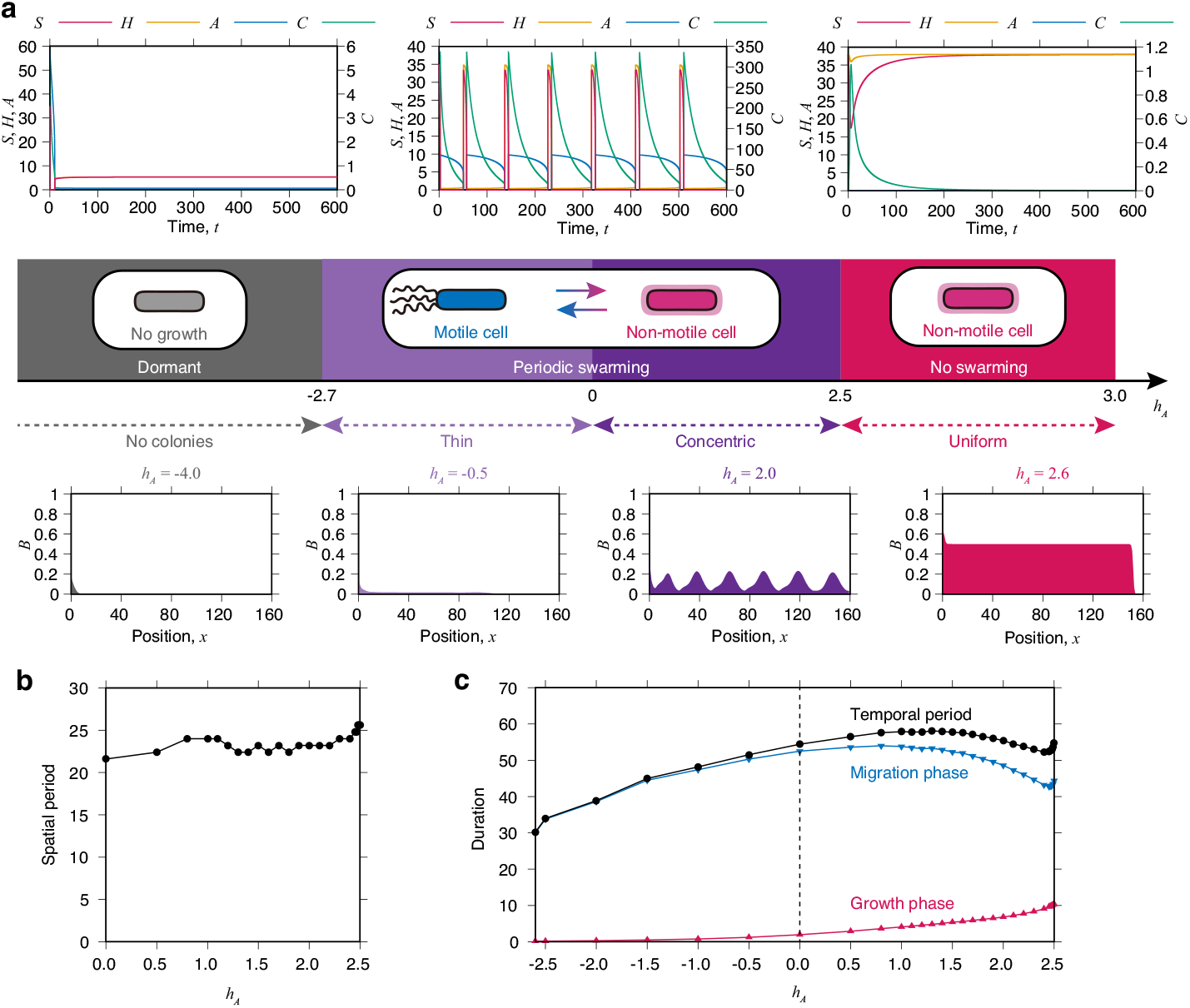
Response of pattern dynamics to changes in *h*_*A*_ (environmental pH) in the simulation. **a**, Changes in cell state at representative points (top) and changes in colony morphology (bottom). **b**, Spatial period (depth of the terrace). **c**, Phase duration: growth phase, migration phase, and their total (migration cycle).

Our simulations revealed distinct patterning regimes based on the value of *h*_*A*_. When *h*_*A*_ > 2.5, the population remains in a non-motile state, corresponding to neutral pH conditions (pH 6.8 or higher). Under these conditions, the colony does not exhibit swarming motility, instead forming spatially uniform patterns driven by cell proliferation and sliding motility (Fig. 5a).

In contrast, within the range of −2.7 < *h*_*A*_ ≤ 2.5, the dominant cell population periodically alternates between motile and matrix-producing states. Specifically, for 0 ≤ *h*_*A*_ ≤ 2.5, stable periodic patterns emerge, which directly corresponds to the experimental stable pH range of 5.3–6.5. While a “thin” pattern appeared for −2.7 < *h*_*A*_ < 0 (a state potentially observable under different conditions such as lower agar concentrations), colonies ceased to grow when *h*_*A*_ < −2.7. This aligns with experimental observations below pH 5.3, where colony formation is inhibited, although the underlying biological mechanisms may involve additional stress responses not fully captured by the current model. Specifically, when lowering pH, cells may cease activity via another mechanism before *h*_*A*_ becomes negative in reality. Furthermore, this diagram depends on various other parameters. By altering factors such as agar concentration and associated motility, or initial nutrient concentration, it may be possible to reproduce responses to diverse environmental changes.

Quantitatively, the periodic patterns observed for 0 ≤ *h*_*A*_ ≤ 2.5 exhibited a nearly constant spatial period (Fig. 5b), which is highly consistent with our experimental data (Fig. 2c,d). Furthermore, the durations of the migration and growth phases remained stable within this range (Fig. 5c), matching the homeostatic behavior observed in vivo (Fig. 2e). We did, however, note a slight divergence near the transition point (*h*_*A*_ = 2.5 vs. pH 6.5). While the simulation successfully reproduced the observed shortening of the migration phase near this boundary, the spatial period in the simulation remained constant (or slightly increased), whereas it shortened in experiments. This discrepancy may suggest that real-world environmental fluctuations near the transition point further reduce swarming duration, leading to the accelerated cycles observed experimentally.

## Discussion

We have developed a multilevel model that generates cell population migration cycles, thereby successfully reproducing the colony formation of *B. subtilis*, which expands periodically to form concentric circles. We sought to elucidate this pattern formation under the assumption that the predominant cell type alternates between motile and non-motile states. Based on extensive molecular biology research, the fundamental gene regulatory network governing the transitions between these two cell types in *B. subtilis* has been well-established. By integrating results from concentric colony culture experiments across various environmental pH levels, we proposed a model for cell-type regulation driven by both environmental conditions and cell density. In this framework, cell density is perceived by the cells through the concentration of the secreted quorum-sensing factor, ComX [37–39]. The mathematical properties of our minimal smooth model also pinpointed the regulatory entry point of environmental pH within the cell-type regulation system [43]. This model allowed us to propose a general control mechanism that selects cell population structures based on environmental stimuli. Notably, the S-shaped equilibrium curve derived from our mathematical analysis provides a robust explanation for the hysteretic control of cell populations forming concentric colonies, and consequently, the mechanism underlying migration cycle generation. Following this concept, we constructed a multilevel model connecting this cell-level state control with tissue-level field equations governing the dynamics of cells, signaling chemicals, and nutrients. Under the simplifying assumption that a single cell type dominates at any given time, this model faithfully reproduced periodically expanding concentric patterns. Furthermore, our analysis demonstrated that environmental pH-induced response can be explained through corresponding parameter variations, providing a consistent framework for understanding changes in both colony morphology diagrams and cell population state control.

The morphology of *B. subtilis* colonies was extensively studied in the 1990s [44, 45], leading to the creation of a morphological diagram where five distinct patterns— Eden cluster-like [46–48], DLA-like [49–54], concentric ring-like, disk-like [44, 55], and DBM-like [56–59]—were identified based on nutrient and agar concentrations. While the concept of discrete cell types was not fully established at that time, our current understanding allows us to reinterpret these patterns: non-motile cells (such as matrix producers) dominate Eden-like and DLA-like colonies, while motile cells predominate in disk-like and DBM-like colonies. Crucially, the concentric colonies focused on in this study are characterized by a periodic switching between these dominant cell types. Among the five patterns, concentric colonies remained a long-standing “unsolved problem” because the mechanism behind this multi-type switching was poorly understood and had not been reproduced by any mathematical model [60]. Our proposed multi-level model is the first to accurately reproduce these concentric patterns, bridging a significant gap in the field.

Historically, the formation of concentric colonies was associated with high uncertainty, occurring only within an extremely narrow range of agar concentrations [22] and showing strong dependence on culture temperature [20]. This suggested that parameters beyond agar and nutrient concentrations were essential for their formation. We hypothesized that the introduction of a stress parameter is necessary to stabilize these patterns and found that environmental pH allows for the reliable formation of concentric colonies. Mapping the morphology diagram within a three-dimensional parameter space enables a more accurate and comprehensive understanding of the entire cell population’s behavior.

Robust biofilms and other microbial populations are complex social structures organized through the coordination of diverse cell types. Therefore, deciphering the mode of state regulation for individual cells is the primary key to understanding and controlling the collective behavior of the population. Higher-order structures in these populations are self-organized in a trans-hierarchical manner, spanning from gene-level and cell-level control to mesoscopic cell distributions, and ultimately to macroscopic morphology, function, and environmental interaction. Our construction of a multilevel model for cell-state control represents a significant step forward in the study of diverse multicellular societies. While our current phenomenological model successfully captures the core dynamics using a simplified assumption of a single dominant cell type, it does not yet predict the fine-scale spatial distribution of coexisting cell types. Future development of new multilevel models capable of describing complex subpopulation structures composed of multiple, coexisting cell types is highly desired.

## Methods

### Experiment and analysis of *B. subtilis* colony cultivation

*B. subtilis* wild-type strain OG-01 (JCM32485) [50, 61] was used in this study. A small inoculum of bacteria (endospores) was incubated overnight (10–16 h) in 5 mL of Luria-Bertani liquid growth medium (LB Broth Base (Lennox L Broth Base), Gibco BRL Life Technologies, cat. no. 12780-052) at 35°C with shaking at 140 rpm. For inoculation onto solid agar media, the bacterial cells were harvested, and the resulting pellet was resuspended in a buffer to a final OD_600_ of 0.5.

The solid medium was prepared using a base solution of 86 mM NaCl and 29 mM K_2_HPO_4_. As a nutrient, 20 g L^−1^ peptone (Bacto Peptone, BD Biosciences, cat. no. 211677) was added to the solution. The pH was then adjusted to a designated value by adding 6 M HCl, followed by the addition of 0.7% agar (Bacto Agar, BD Biosciences, cat. no. 214010). After autoclaving at 121°C for 15 min, 20 mL of the solution was poured into each plastic Petri dish. The dishes were maintained at room temperature overnight and subsequently dried at 50°C for 90 min.

The pre-cultured bacterial suspension was inoculated using a sterilized needle at the center of the prepared solid medium surface. The dishes were then incubated in a humidified box at 35°C for the designated period. Macroscopic images of the colonies were obtained via transmitted light using a flatbed scanner (GT-X980, Epson). For higher-resolution observations, cellular and cluster-level morphology was examined by phase-contrast microscopy using an inverted microscope (IX83, Olympus) equipped with a digital camera (DP80, Olympus).

The raw image data (Extended Data Fig. 1a) were converted into cell density distributions (Extended Data Fig. 1b), which were then used to reconstructed 3D structures (Extended Data Fig. 1c) and analyze cross-sections (Extended Data Fig. 1d). By performing these quantitative analyses in a time series, we were able to evaluate not only the two-dimensional cell density (colony thickness) but also the dynamics of cell proliferation and migration (Fig. 1).

### Diversity of *B. subtilis* colony patterns selected in response to the environment

Although the requirement for agar concentrations of approximately 0.7% and high-nutrient conditions in solid media was previously recognized [20, 22], the precise conditions for forming concentric *B. subtilis* colonies had not been clearly established. Specifically, the agar concentration must be within a narrow range of 0.65–0.75%. Furthermore, the preferred incubation temperature is 25–35°C —lower than the standard 35–37°C [20]—and extended pre-drying of the medium surface is often necessary [22]. These conditions are further influenced by temperature and surface moisture, in addition to agar and nutrient concentrations. To address this complexity, we introduced environmental pH as a novel parameter to clarify the growth requirements of *B. subtilis*.

In most colony types other than concentric ones, the dominant cell population consists of either motile cells or matrix producers. Colonies driven by swarming motile cells typically develop at agar concentrations of 0.4–0.7%. These are categorized into two types based on nutrient concentration: disk-like (high nutrient) [44, 55] and dense branching morphology (DBM)-like (low nutrient) [56–59]. Conversely, slowly expanding colonies dominated by matrix producers form on stiff media with agar concentrations of 0.7% or higher. These are also classified into two types depending on nutrient availability [50, 51]: Eden cluster-like (high nutrient) [46–48] and diffusionlimited-aggregation (DLA)-like (low nutrient) [49–52]. While these five types form the basis of general classification, detailed examinations have shown that even on stiff media (e.g., 1% agar), a small number of motile cells can coexist on the surface, influencing macroscopic morphology and expansion rates [62]. Notably, a crater-like pattern, where cells aggregate at the colony periphery, can form when the environmental pH is shifted only slightly below the intracellular pH of 7.4. This phenomenon is driven by chemotaxis toward nutrients ^―^a behavior that does not occur during swarming [28]. On the other hand, at lower agar concentrations (0.5%), rapid colony expansion via swarming is observed, as abundant surface moisture reduces the energy required for migration. However, under specific conditions, such as an optimal pH of 7.4, matrix producers may become the dominant cell type instead of motile cells, resulting in lacework-patterned colonies [24]. Thus, environmental pH serves as a critical stress parameter influencing the cell-type selection between motile cells and matrix producers, even outside the context of concentric colonies.

### Microscopic observation of phase switching in concentric colonies

Using phase-contrast microscopy, we investigated the formation of concentric colonies and the spatio-temporal dynamics of cell states. The termination of the matrix-producer-dominant growth phase begins with the emergence of motile cells just inside the colony boundary (Extended Data Fig. 2a). These motile cells continue to proliferate, forming pools where active swarming occurs. The surrounding matrix-producing cells are partially dissociated by the action of autolysins [63–66], and the increasing mechanical pressure from the proliferating motile cells eventually leads to multiple breaches along the colony border.

From these ruptured points on the outer edge of the colony, motile cells flow outward to form branches (Extended Data Fig. 2b,c, Supplementary Video S1). The subsequent migration phase, dominated by motile cells, is characterized by the rapid radial expansion of these branches via swarming (Extended Data Fig. 2d, Supplementary Video S2). As the number of breaches near the branch roots increases, the gaps between branches are filled by outflowing cells from the interior (Extended Data Fig. 2e,f, Supplementary Videos S3, S4). The migration phase concludes when the outward growth of the branches ceases (Extended Data Fig. 2g). Immediately thereafter, the swarming region recedes from the tips of the branches toward the inner roots (Extended Data Fig. 2g–i). Once the swarming activity completely disappears, the population reverts to the growth phase, during which cells remain stationary while continuing to proliferate, leading to an increase in local cell density. When the cell concentration exceeds a critical threshold, motile cells reappear, triggering the next transition to the migration phase (Extended Data Fig. 2a). This sequence constitutes one complete round of the migration cycle in the *B. subtilis* cell population.

### Numerical simulation procedures and model implementation

Numerical calculations for the multilevel model ((1)–(2) with 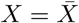) were performed using the Euler method. The colony radius is defined as the largest value of *x* ∈ Ω for which 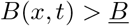, where 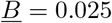 is a predetermined threshold constant.

To represent the distinct modes of bacterial movement, the motility term *D*_*B*_ is efined by a switch between motile and non-motile states:

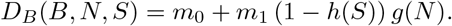

ere, *m*_0_ represents sliding motility, which involves passive movement on the medium rface by matrix producers, and *m*_1_ represents the flagellar motility of motile cells. he transition to the motile cell state is determined by the master regulator *S* exceedg a threshold *S*_mot_. This switching is modeled using a Heaviside function *h* = *h*(*S*) ith a threshold *S*_mot_:

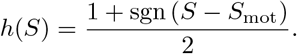

Alternatively, a Hill function could be employed: 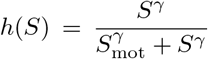, where *γ* is a sensitivity exponent. The dependence of motility on nutrient concentration *N* follows a Hill function 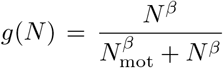 with an exponent *β* and a constant *N*_mot_. The qualitative results of the model are robust and do not fundamentally depend on the specific choice of these switching functions.

Other biological processes, including cell proliferation *p* and the secretion of the signaling molecule *s*, are likewise state-dependent. We assume that matrix producers, which dominate the growth phase, exhibit higher rates of proliferation and *X* secretion mpared to motile cells: *p*(*B, N, S*) = *p*_1_*BNh*_1_(*S*), *s*(*B, N, S*) = *s*_1_*Bg*_1_(*N*)*h*_1_(*S*),where *p*_1_ and *s*_1_ are positive constants, 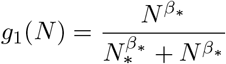with an exponent *β*_*_ and a constant *N*_*_. Similarly to the motility, we adopt the Heaviside function with a thresh-old *S* for representing the switching of cell type properties: 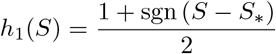 for a constant *S*_*_. Nutrient consumption is assumed to occur at a common rate for both cell types for simplicity: *c*(*B, N, S*) = *c*_1_*BN*, where *c*_1_ is a positive constant.

Calculations were conducted on a one-dimensional spatial domain with Neumann oundary conditions at both ends:

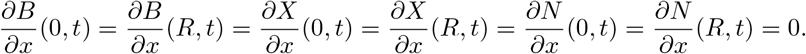

The initial nutrient concentration is spatially uniform: *N* (*x*, 0) = *N*_0_ for a constant *N*_0_. The initial concentration of the signaling substance *X* is set to 0: *X*(*x*, 0) = *X*_0_ = 0. Bacteria were inoculated at the left boundary (*x* = 0), representing the center of the colony in a cross-sectional view (i.e., in the simulation, we are viewing the right half of the colony’s cross-section):

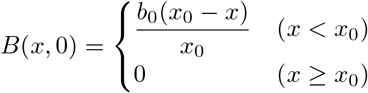

where *x*_0_ denotes the width of the inoculation region and *b*_0_ represents the inoculation amount (maximum initial cell density). The complete set of parameters used in the simulations is provided in Extended Data Table 1.

## Supporting information

Legends for Supplementary Videos S1-S4

## Code availability

All code is available via GitHub at https://github.com/sohei-tasaki/sigrep-SHAC-BXN.

## Acknowledgements

This work was supported by JSPS KAKENHI, Grant Number 23K03225 (M.N.), 26K06907 (M.N.), 23K03176 (I.T.), 19K03645 (S.T.), 23K03208 (S.T.), and MEXT KAKENHI, Grant Number 17H06327 (S.T.).

---

**Extended Data Table 1:**
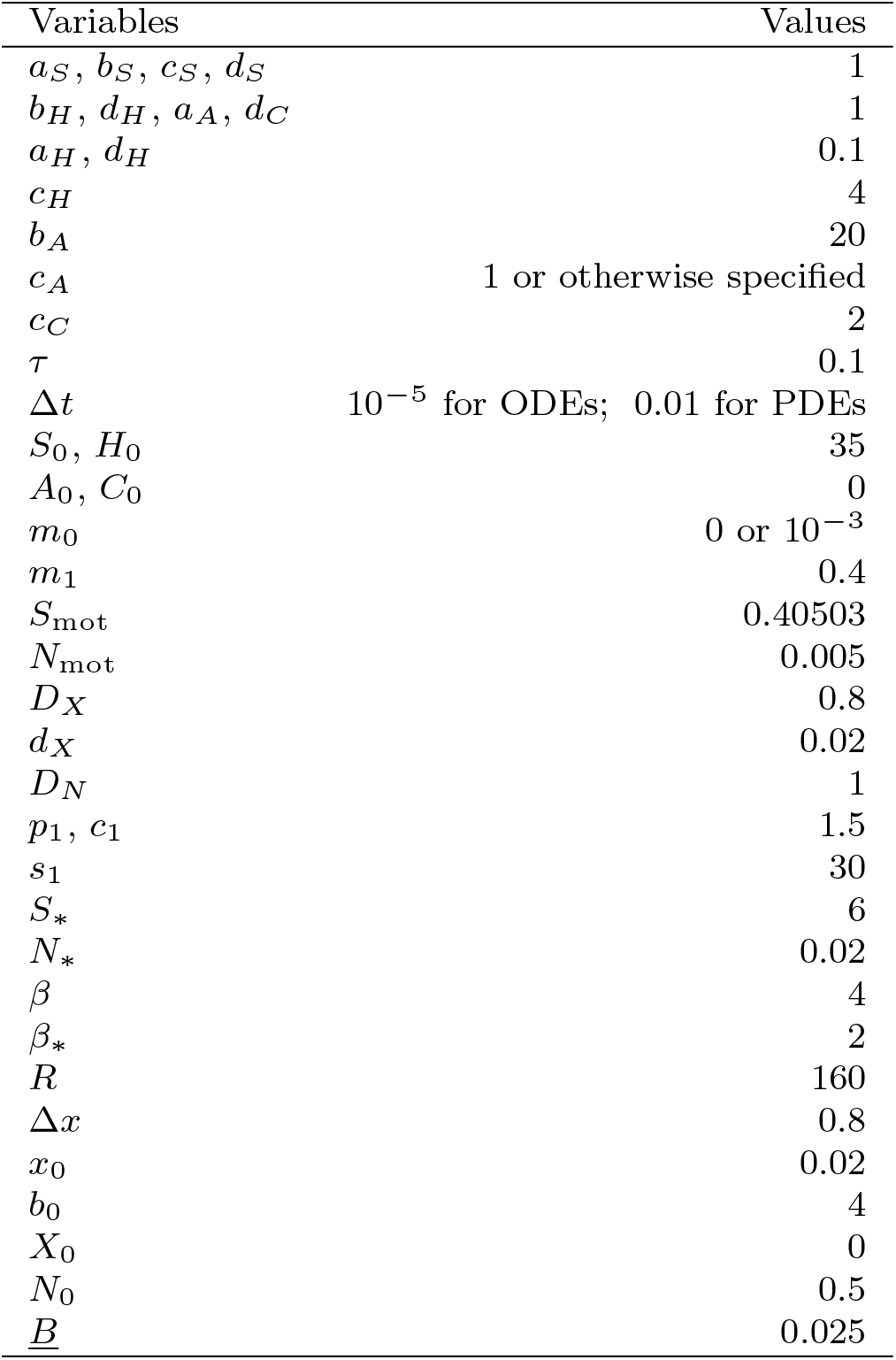
The parameter values used in our simulations.

| Variables | Values |
| --- | --- |
| $a_S, b_S, c_S, d_S$ | 1 |
| $b_H, d_H, a_A, d_C$ | 1 |
| $a_H, d_H$ | 0.1 |
| $c_H$ | 4 |
| $b_A$ | 20 |
| $c_A$ | 1 or otherwise specified |
| $c_C$ | 2 |
| $\tau$ | 0.1 |
| $\Delta t$ | $10^{-5}$ for ODEs; 0.01 for PDEs |
| $S_0, H_0$ | 35 |
| $A_0, C_0$ | 0 |
| $m_0$ | 0 or $10^{-3}$ |
| $m_1$ | 0.4 |
| $S_{\text{mot}}$ | 0.40503 |
| $N_{\text{mot}}$ | 0.005 |
| $D_X$ | 0.8 |
| $d_X$ | 0.02 |
| $D_N$ | 1 |
| $p_1, c_1$ | 1.5 |
| $s_1$ | 30 |
| $S_*$ | 6 |
| $N_*$ | 0.02 |
| $\beta$ | 4 |
| $\beta_*$ | 2 |
| $R$ | 160 |
| $\Delta x$ | 0.8 |
| $x_0$ | 0.02 |
| $b_0$ | 4 |
| $X_0$ | 0 |
| $N_0$ | 0.5 |
| $\underline{B}$ | 0.025 |

**Extended Data Fig. 1:**
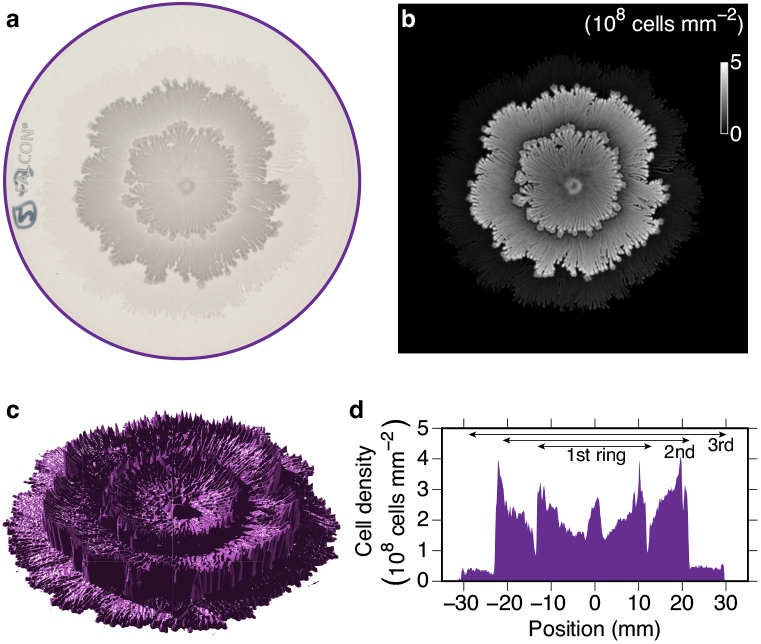
Data conversion to a cell density field and its visualization. **a**, Raw image of transmitted light obtained by a flat scanner. The diameter of the dish is 88 mm. **b**, Two-dimensional cell density. **c**, Three-dimensional reconstruction of the colony. **d**, Cross-sectional view.

**Extended Data Fig. 2:**
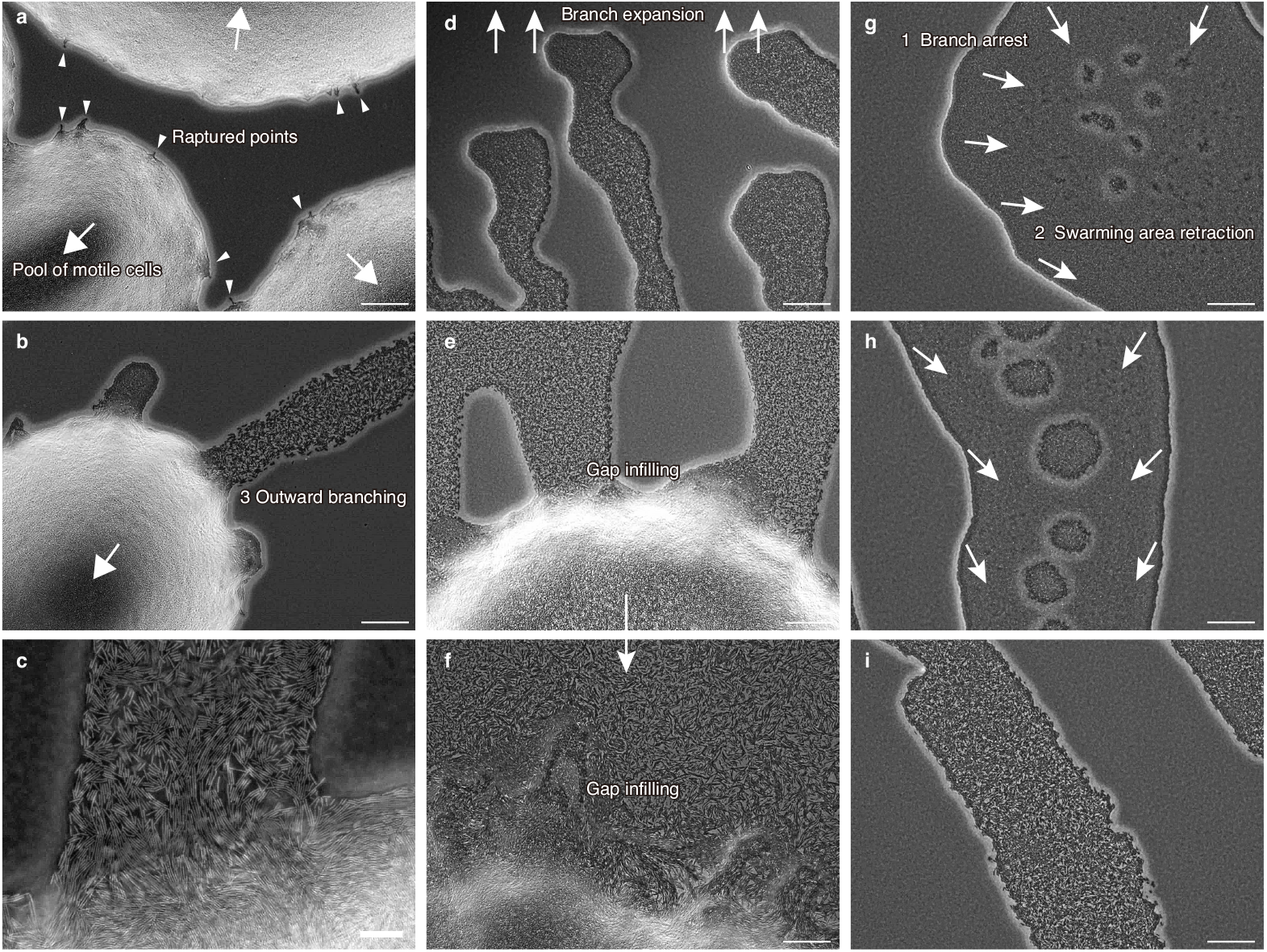
Microscopic images of phase transition during colony expansion. **a–c**, Transition from the growth phase to the migration phase. Motile cells appear slightly inside colony boundaries (large arrows in **a** and **b**). The number of motile cells increased to form bacterial pools where cells organize to undergo swarming collective motion. The motile cells then destroy the boundary walls (small triangles in **a**). Cells flow out from some of the outermost raptured points on the colony boundaries to create rapidly expanding branches (**b** and **c**). **d**–**f** The migration phase. Numerous dense branches grow outward in **d**. Gradual collapse of the colony boundaries at the roots of branches, and the space around the roots is filled with the outflowing cells (**e** and **f**). **g**–**i** Transition from the migration phase to the growth phase. First, the branches stop growing in **g**. Then, the swarming regions composed of motile cells (small bubble-like regions in **g** and **h**; whole branch region in **i**) retract from the branch tips toward the colony interior (**g, h** to **i**; **g** is the outermost boundary, **i** is the innermost). Scale bars, 100 *µ*m (except for **c**), 10 *µ*m in **c**.

**Extended Data Fig. 3:**
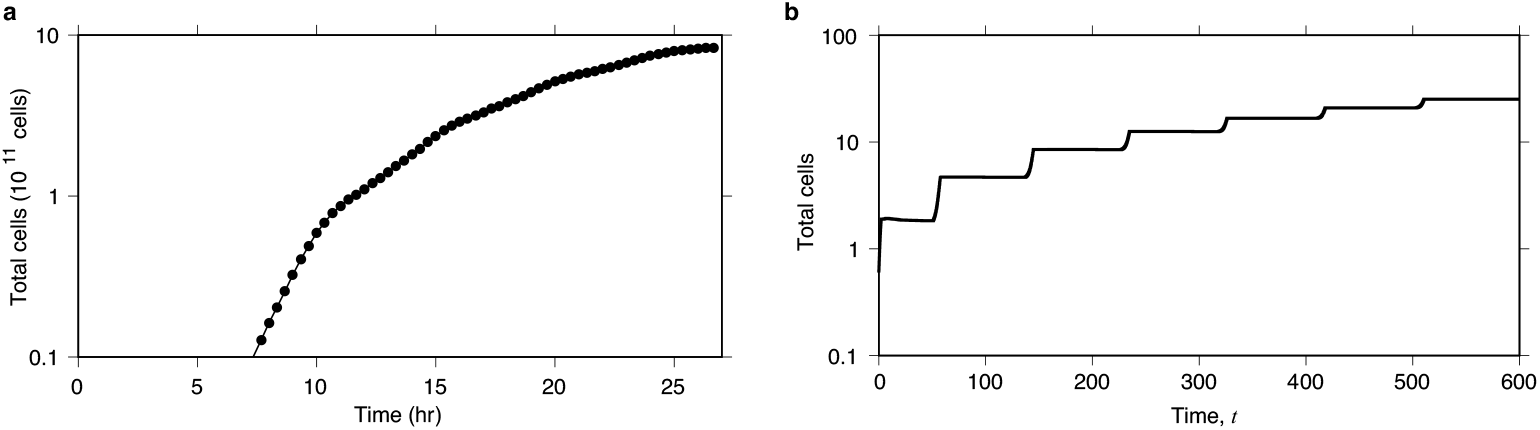
Time series of total cell number during concentric colony growth. **a**, Experiment. **b**, Simulation.

**Extended Data Fig. 4:**
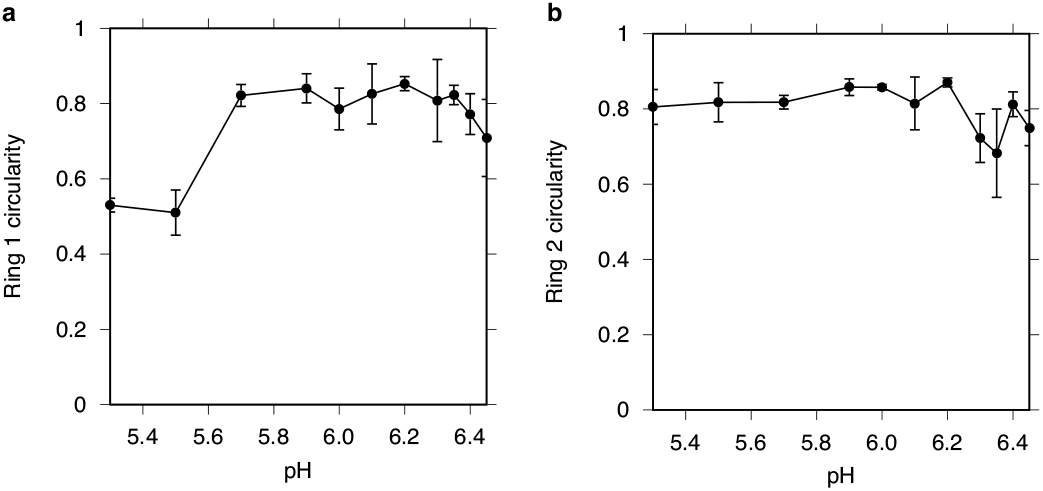
Circularity 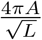 of the rings in concentric colonies in response to environmental pH fluctuations, where *A* is the area, *L* is the circumference). **a**,Circularity of first ring. **b**, Circularity of second ring.

