## Supplementary material for "Multilevel modeling of cell population migration cycles and response to environmental pH in *Bacillus subtilis*": Legends for Supplementary Videos S1-S4

Running title: Multilevel modeling of *B. subtilis* migration cycles

### Legends for Supplementary Videos S1-S4

**Supplementary Video S1.** Cells flow out from some of the outermost breach points at the colony’s boundary, forming rapidly outward-growing branches.

**Supplementary Video S2.** Numerous branches composed of swarming cells extend outward.

**Supplementary Video S3.** Gradual collapse of the colony boundaries at the roots of branches. The space around the roots is filled with the outflowing cells (Supplementary Video S4 shows a later stage).

**Supplementary Video S4.** Gradual collapse of the colony boundaries at the roots of branches. The space around the roots is filled with the outflowing cells (Supplementary Video S3 shows an earlier stage).
